# ARL15 modulates magnesium homeostasis through N-glycosylation of CNNMs

**DOI:** 10.1101/2020.09.09.289835

**Authors:** Yevgen Zolotarov, Chao Ma, Irene González-Recio, Serge Hardy, Gijs Franken, Noriko Uetani, Femke Latta, Elie Kostantin, Jonathan Boulais, Marie-Pier Thibault, Jean-François Côté, Irene Díaz Moreno, Antonio Díaz Quintana, Joost G.J. Hoenderop, Luis Alfonso Martínez-Cruz, Michel L. Tremblay, Jeroen H.F. de Baaij

**Affiliations:** Rosalind and Morris Goodman Cancer Research Centre, McGill University, Montréal, QC, Canada H3A 1A3; Department of Biochemistry, McGill University, Montréal, QC, Canada H3G 1Y6; Department of Physiology, Radboud Institute for Molecular Life Sciences, Radboud university medical center, 6500HB, Nijmegen, The Netherlands; Liver Disease Laboratory, Center for Cooperative Research in Biosciences (CIC bioGUNE), Basque Research and Technology Alliance (BRTA), Bizkaia Technology Park, Building 801A, 48160 Derio, Spain; Montreal Clinical Research Institute (IRCM), Montréal, QC, Canada H2W 1R7; Institute for Chemical Research (IIQ), Scientific Research Centre “Isla de la Cartuja” (cicCartuja), University of Seville - CSIC, Avda. Américo Vespucio 49, Seville, 41092, Spain

**Keywords:** CNNM2, CNNM3, Glycosylation, Magnesium transport, Protein-protein interaction

## Abstract

Cyclin M (CNNM1-4) proteins maintain cellular and body magnesium (Mg^2+^) homeostasis. Using various biochemical approaches, we have identified members of the CNNM family as direct interacting partners of ADP-ribosylation factor-like protein 15 (ARL15), a small GTP-binding protein. ARL15 interacts with CNNMs at their carboxyl-terminal conserved cystathionine-β-synthase (CBS) domains. *In silico* modeling of the interaction using the reported structures of both CNNM2 and ARL15 supports that the small GTPase specifically binds the CBS1 domain. Immunocytochemical experiments demonstrate that CNNM2 and ARL15 co-localize in the kidney, with both proteins showing subcellular localization in the Golgi-apparatus. Most importantly, we found that ARL15 is required for forming complex N-glycosylation of CNNMs. Overexpression of ARL15 promotes complex N-glycosylation of CNNM3. Mg^2+^ uptake experiments with a stable isotope demonstrate that there is a significant increase of ^25^Mg^2+^ uptake upon knockdown of ARL15 in multiple kidney cancer cell lines. Altogether, our results establish ARL15 as a novel negative regulator of Mg^2+^ transport by promoting the complex N-glycosylation of CNNMs.

## INTRODUCTION

Mg^2+^ is an essential cation for all living organisms. Mg^2+^ deficiency, therefore, causes organ dysfunction and is associated with a wide range of human diseases [1]. As co-factor of ATP, Mg^2+^ is involved in over 600 enzymatic reactions and as such it plays important roles in a plethora of biological mechanisms including DNA transcription, protein synthesis and energy metabolism [1]. For example, decreased intracellular Mg^2+^ levels suppress cell cycle progression and are consequently associated with cell growth impairment [2]. Recent studies therefore have focused on intracellular Mg^2+^ concentration as a central mechanism for cellular metabolism and proliferation [3].

Among the various mechanisms of Mg^2+^ sensing and transport, proteins of the Cyclin M (CNNM) family were shown to play an important role in the intracellular sensing of the Mg^2+^ availability [4][5]. The CNNM family has 4 members in mammalians, which are highly conserved but have different tissue distribution[4][6][3][5]. CNNM1, CNNM2 and CNNM4 have highest expression levels in brain, kidney and intestine, respectively whereas CNNM3 shows a ubiquitous expression pattern [7]. Genetic defects of CNNMs have been demonstrated to be causative for disease. Mutations in *CNNM2* are causative for a multi-organ syndrome comprising intellectual disability, seizures and renal Mg^2+^ wasting [OMIM: #616418] [8]. Patients with CNNM4 mutations suffer from Jalili syndrome, consisting of cone-rod dystrophy and amelogenesis imperfecta [OMIM: #217080] [9].

CNNMs interact with phosphatases of the regenerating liver (PRL1-3) via the cystathionine-β-synthase (CBS) domain of CNNMs [10][11]. CNNMs and PRLs have been associated with progression of breast and colon cancers [11][12]. Although the exact function of CNNM proteins is under debate [13][14], it has been shown that engineered and naturally occurring mutations in CNNMs result in decreased Mg^2+^ transport in both cell and animal models [8][11].

Recently, a large GWAS identified the ADP-ribosylation factor-like protein 15 (*ARL15*) locus to be associated with urinary Mg^2+^ excretion [15]. ARL15 is structurally similar to Ras-related GTP-binding proteins, which regulate intracellular vesicle trafficking [16][17]. Within the kidney, ARL15 is highly expressed in the thick ascending limb (TAL) and distal convoluted tubule (DCT), like CNNM2, where Mg^2+^ reabsorption is tightly regulated [15]. However, the exact function of ARL15 and the mechanism by which ARL15 regulates renal Mg^2+^ handling are still unknown.

In this study, we identified CNNMs as direct interacting partners of ARL15 by binding to the CBS1 domain of CNNM2. We demonstrate that ARL15 is a negative regulator of cellular Mg^2+^ transport through its modulation of CNNMs N-glycosylation and the direct correlation of its expression to levels of intracellular Mg^2+^.

## RESULTS

### ARL15 is a new binding partner of CNNMs

To identify protein interactions of CNNMs, pull-down of mouse CNNM2 carboxyl (C)-terminal region was performed followed by mass spectrometry (Fig. 1A). Given that CNNM2 is known to regulate Mg^2+^ reabsorption in the DCT segment of the nephron, DCT-enriched kidney lysates were used. Twenty-four interacting partners of CNNM2 were detected, including ARL15 (Fig. 1B). Since ARL15 has been shown to be involved in urinary Mg^2+^ excretion [15], we decided to further explore this particular interacting partner. To confirm the interaction between ARL15 and CNNMs, BioID of ARL15 was performed and we identified 221 proteins as potential interacting partners of ARL15 and CNNM2, CNNM3 and CNNM4 were among them (Supp. Fig. 1). GO biological process overrepresentation analysis indicated that Mg^2+^ homeostasis is one of the most highly enriched processes (Fig. 1C). Immunoprecipitation demonstrated that CNNM3 and ARL15 interact endogenously (Fig. 1D). To further confirm this association, co-immunoprecipitation assays were performed with overexpressed ARL15 and the four members of the CNNM family. ARL15 binding was observed to a similar extent for each CNNM protein (Fig. 1E). Endogenous PRL-1 and PRL-2, which are well described interacting partners of CNNMs that regulate Mg^2+^ flux [12][11], also co-precipitated with CNNMs and ARL15 (Fig. 1E). The above data indicates that ARL15 is a novel interacting partner of the four members of the CNNM family of Mg^2+^ modulators.

**Fig. 1.**
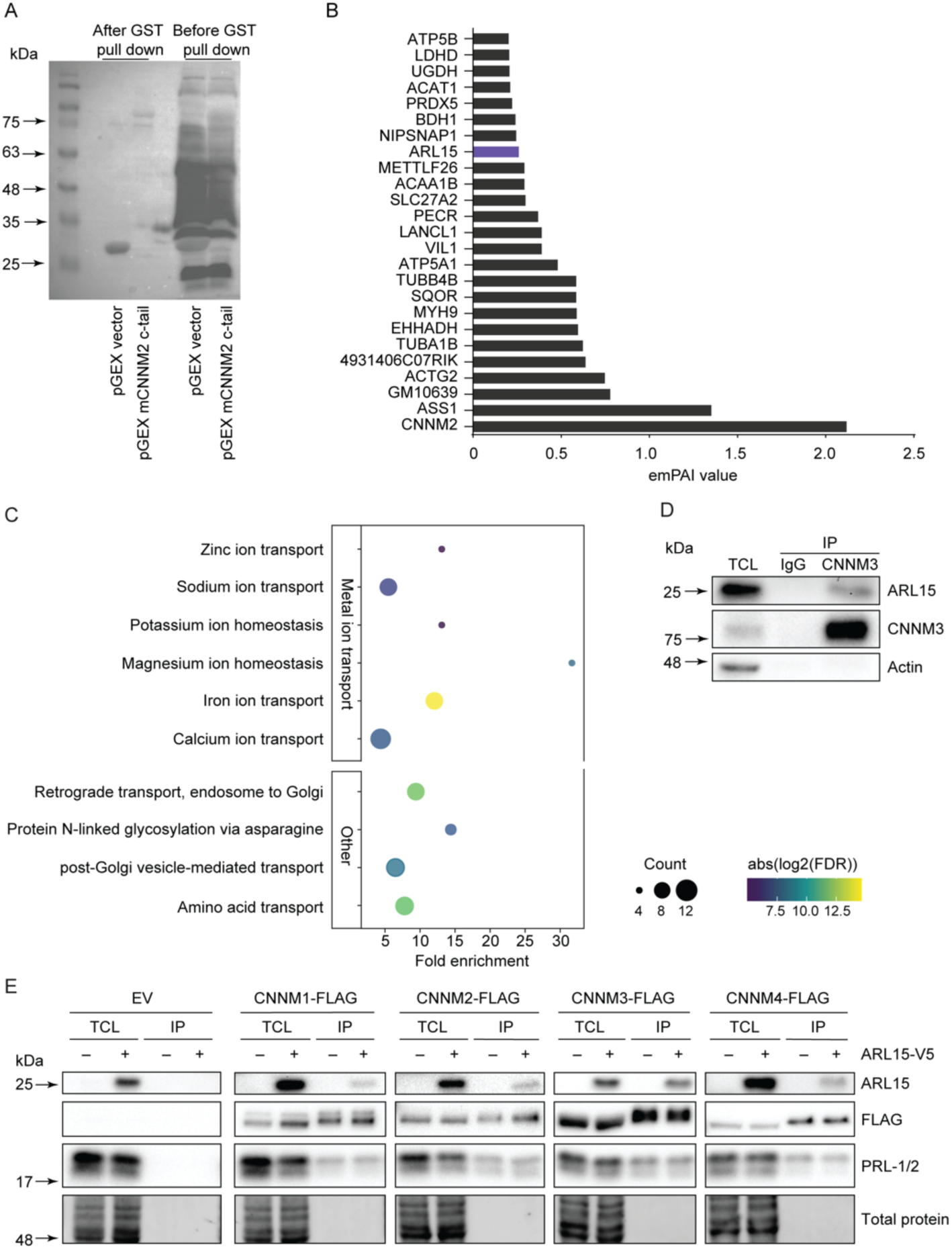
ARL15 interacts with CNNMs. A. Immunoblot of GST-mCNNM2 c-tail pGex and empty pGex was used as control. The C-tail of mCNNM2 is purified and bind to glutathione beads. The DCT enriched fraction from 8 parvalbumin GFP mice were added to the bead. **B**. List of binding proteins to mCnnm2 C-tail (emPAI value). **C**. Representative pathways related to metal ion transport and Golgi-trafficking identified using gene ontology biological process overrepresentation analysis with ARL15 interacting partners. **D**. Endogenous CNNM3 was immunoprecipitated from HEK293 lysates and endogenous ARL15 co-immunoprecipitated with it. IgG antibody of the same subclass as CNNM3 antibody was used as negative control. **E**. Anti-FLAG beads were used to immunoprecipitate overexpressed FLAG-tagged CNNM1-4 proteins. The blot shows that ARL15 interacts with CNNM1-4, as well as endogenous PRL-1 and PRL-2. emPAI, Exponentially modified protein abundance index; TCL, Total cell lysate; IP, Immunoprecipitation; GST, Glutathione-S-transferase; ARL5, ADP ribosylation factor like GTPase 15; CNNM, Cyclin M.

### ARL15 interacts with CNNM2 cytoplasmic region

The topology for CNNMs consists of a signal peptide, three transmembrane domains (TM) and an intracellular C-terminus containing the Bateman module (which in turn is formed by two consecutive CBS domains, CBS1 and CBS2) and a cyclic nucleotide monophosphate-binding homology domain (CNBH) domain) [7]. To determine which domain of CNNM2 is important for the interaction with ARL15, truncated CNNM2 proteins were co-immunoprecipitated with ARL15 (Fig. 2A). The presence of the CBS domains is essential for the binding of ARL15 to CNNM2. The interaction of the proteins decreased by 36% when the CBS2 domain is removed (Fig. 2B and 2C). No interaction with the transmembrane domain was observed. The interaction between ARL15 and the Bateman module of CNNM2 was confirmed by gel filtration chromatography (Fig. 2D and 2E). Pure ARL15 and CNNM2_BAT_ eluted as homodimers when they were independently loaded into a size exclusion chromatography column (*V*_e(ARL15)_≈15.02 mL, *M*_exp(ARL15)_≈ 36.03kDa; *V*_e(CNNM2BAT)_≈14.67mL, *M*_exp(CNNM2BAT)_≈ 43.98 kDa, respectively, where *V*_e_ indicates the elution volume and *M*_exp_ indicates the experimental molecular weight). The corresponding CNNM2_BAT_·ARL15 complex (peak marked with *black asterisk*) eluted as a heterotetramer 2xARL15+2xCNNM2_BAT_ (*V*_e_≈13.77 mL, *M*_th_= 74.58 kDa, *M*_exp_≈ 73.41 kDa).

**Fig. 2.**
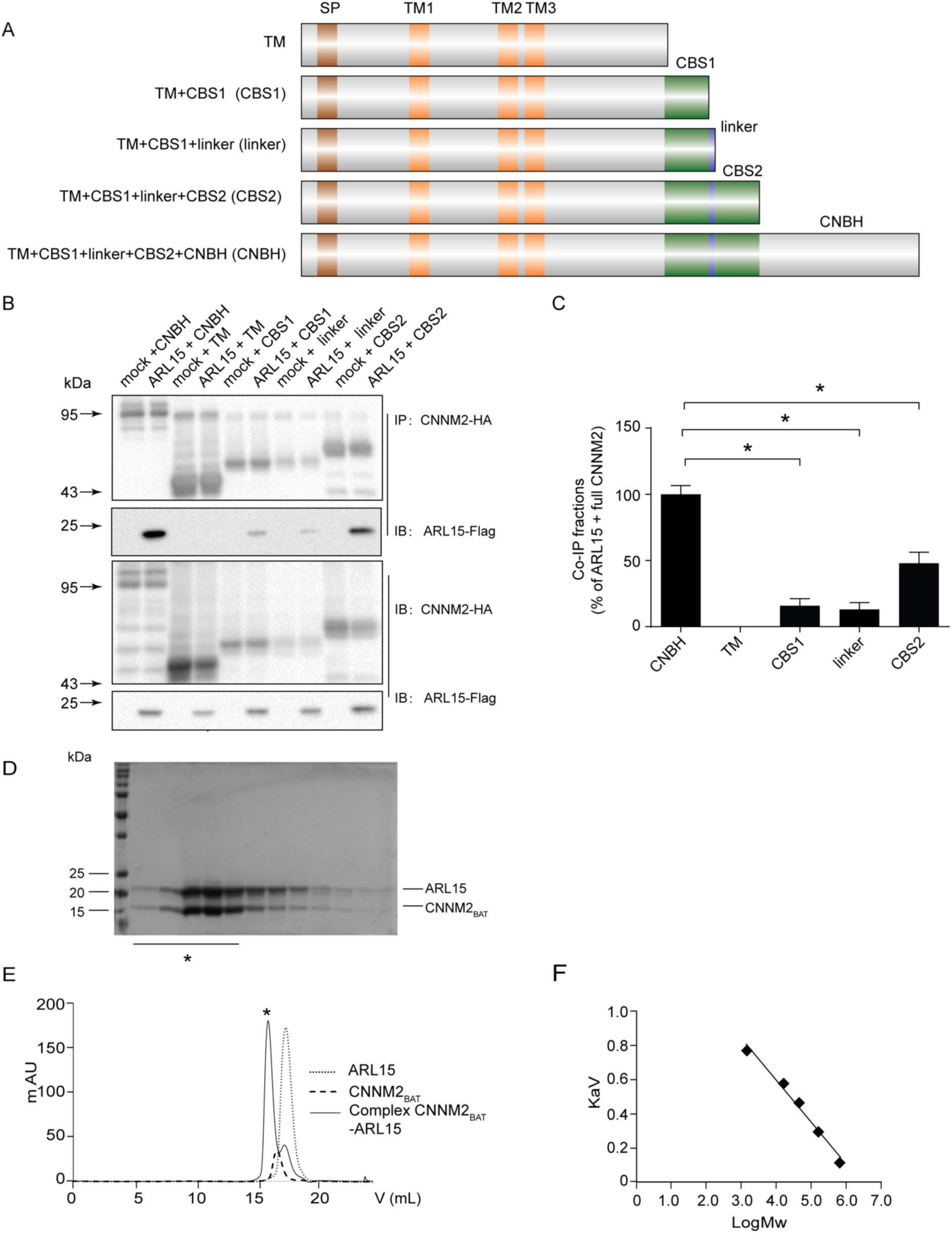
ARL15 interacts with CNNM2 cytoplasmic region. **A**. Schematic overview of CNNM2 constructs including the predicted protein domains and truncations. **B**. HEK293 cells were co-transfected with truncated CNNM2-HA and ARL15-Flag. The upper two blots show the detection of the Flag-tagged proteins in anti-HA precipitated cell lysates. The lower two blots show input controls of HA-tagged and Flag-tagged proteins respectively. **C**. Quantification of ARL15/CNNM2 binding between the different truncated CNNM2 proteins. Results are the mean ± SEM of 3 independent experiments. * indicate significant differences compared to CNNM2 + ARL15 transfected cells (P<0.05). SP, Signal peptide; TM, Transmembrane; CBS, Cytosolic cystathionine β-synthase; CNBH, Cyclic nucleotide monophosphate-binding homology domain; ARL5, ADP ribosylation factor like GTPase 15; CNNM, Cyclin M. D. SDS-polyacrylamide gel run with the isolated fractions of the CNNM2_BAT_·ARL15 complex. Bands correspond to the isolated peak (marked with *black asterisk* in E) of the chromatographic run of the CNNM2_BAT_·ARL15 complex (*M*_th(ARL15monomer)_=19.57 kDa and *M*_th(CNNM2BATmonomer)_= 17.72 kDa, where *M*_th_ indicates the theoretical molecular weight of each subunit). E, *left*. Gel filtration of the CNNM2_BAT_.ARL15 complex and of its individual components. Bands correspond to the peak (marked with *black asterisk*) of the chromatographic run of the complex CNNM2_BAT_·ARL15 (*M*_th(ARL15monomer)_=19.57 kDa and *M*_th(CNNM2BATmonomer)_= 17.72 kDa, where *M*_th_ indicates the theoretical molecular weight of each subunit). E, *right*. The calibration trendline calculated from protein standards is *y*= −0.2451*x* + 1.5726 (*y* = *K*_av_; *x* = log*M*).

To determine whether the interaction between ARL15 and CNNMs is dependent on Mg^2+^, ARL15 and CNNM were immunoprecipitated in the presence or absence of Mg^2+^. Results showed that the presence of 1 mM Mg^2+^ in the lysis buffer does not affect the association between ARL15 and CNNM2 compared to the condition of the lysis buffer without Mg^2+^ (Supp. Fig. 2A). In line with this, ARL15 and CNNM3 binding was similar in both conditions (Supp. Fig. 2B).

### Docking models of the CNNM-ARL15 interaction

To assess the role of the various cytoplasmic CNNM2 cytosolic domains as targets, ARL15 docking computations were carried out. The coordinates of the flexible linker joining the Bateman module with the CNBH domain were modelled onto the available X-ray diffraction data of the CNNM2 cytosolic domains. Figure 3 displays the domain arrangement of CNNM2 cytosolic domains (Fig. 3A) as well as the electrostatic potential on their surface (Fig. 3B). Notably, the linkers joining the Bateman domain to the CNBH display a stretch of basic residues. Best score solutions displayed ARL15 interacting with both CNNM2 monomers (Supp. Movie 1). ARL15 was partly inserted into the cleft between the Bateman and the CNBH domains (Fig. 3C-D). In each of the two symmetric solutions, the total buried solvent accessible surface per partner was 2,175.7 Å^2^—1,830.8 Å^2^ (932.6 Å^2^, from CBS1, 262.3 Å^2^ from the linker and 344.9 Å^2^ from the CNBH domain) with one of the two CNNM2 monomers and 344.9 Å^2^ (190.6 Å^2^ from CBS1* and 154.3 Å^2^ from CNBH*) for the interaction with the second CNNM2 monomer. Such values are typical of rather stable complexes, and in the upper limit of non-obligate interactions [18]. In addition, the complex displays complementarity of charges at the rim of the interface, as shown in Fig. 3D. As pointed out, the CBS1 domains from complementary Bateman modules in the CNNM2 dimer contacted with ARL15, in particular that of the first monomer with a surface region defined between α−helix A, the β1-β2 hairpin loop and α-helix H1 in one of them, but also contacting the second monomer. Figure 3C shows the lowest energy solution found in different simulations. Other solutions with low energies corresponded to the hydrophobic surface of H0 helix pair, which interacts with the lipid bilayer, and spots of the CNBH domains in which the absence of coordinates from the nearby unstructured regions disclosed an artifactual interaction site, otherwise concealed. Notably, Brownian Dynamics computations (Supp. Fig. 3A, Supp. Table1) yielded results very similar to those displayed herein (Supp. Fig. 3). Thus, the interaction model herein is compatible with a 1:1 CNNM2:ARL15 stoichiometry. Other sets of solutions targeted the same region, but displayed different orientations of ARL15 towards CNNM2 surface, and were not found in all simulations. The result in Fig. 3, however, was hardly dependent on the modelled conformation of the linker residues 525 to 546 (Fig. 3D). Nevertheless, in most simulations several positive residues of the linker mapped near the interface. As CNNM3 has been reported to bind ARL15 with an affinity higher than that for CNNM2, HEX docking simulations were also performed using a CNNM3 model (Supp. Fig. 3). Notably, ARL15 targeted the same site in CNNM3 as does in CNNM2, though the orientation differs by a slight rotation of ARL15.

**Fig. 3.**
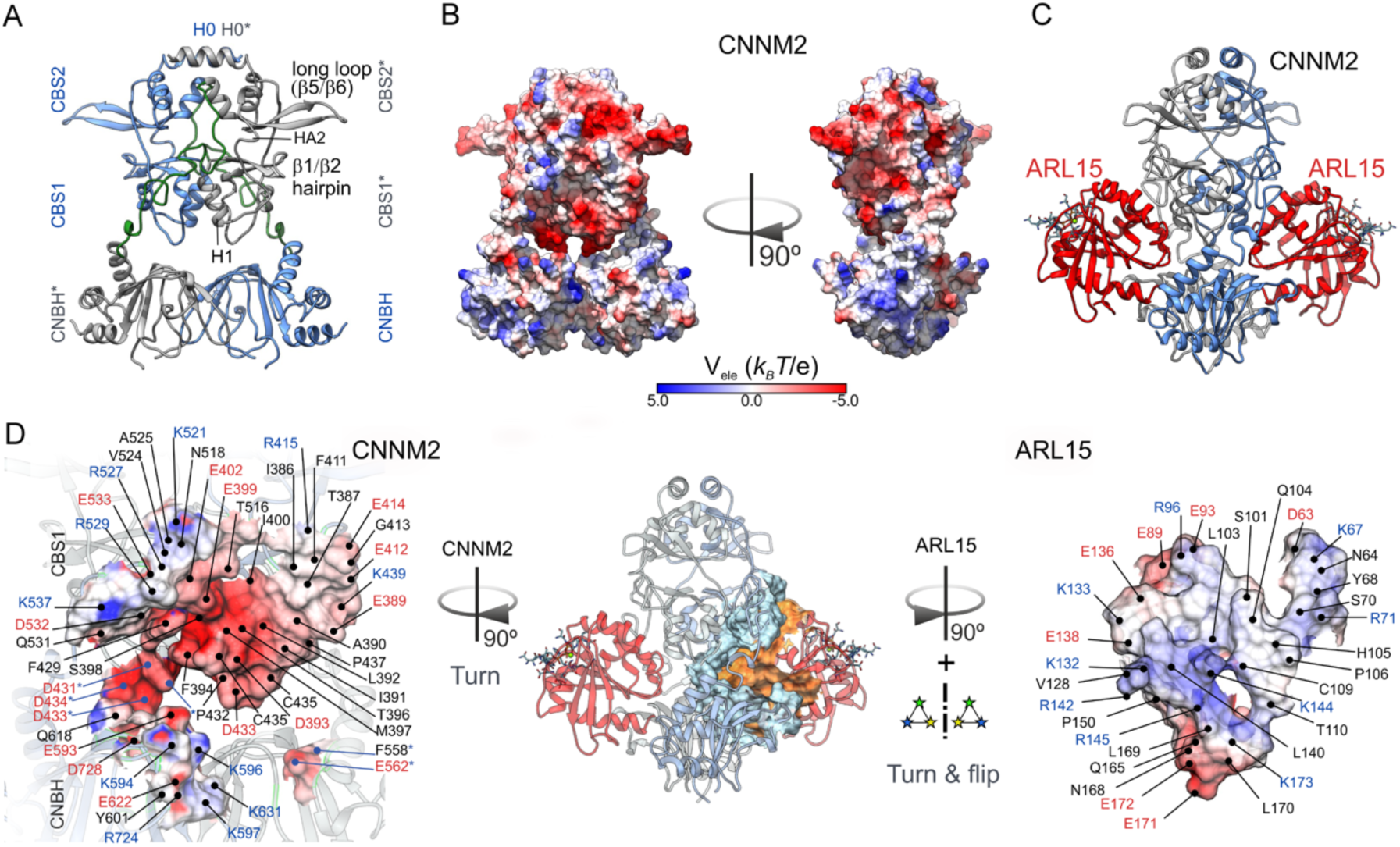
Structural model of the interaction between the CNNM2 cytosolic domains and ARL15. **A**. Richardson (ribbon) diagram of the cytosolic CNNM2 dimer domains. Each monomer is in a different color, and the model of flexible linker joining the Bateman domains to the CNBH ones are in light green. **B**. Electrostatic potentials at the surface of CNNM2 cytosolic domains. Electrostatic grids were generated with APBS 3.0 (Baker *et al*, 2001) at 150 mM ionic strength. **C**. Best solution of ARL15 (red ribbons) docking to CNNM2 computations. **D**. Dissection of the interface between ARL15 and CNNM2 (middle) illustrating both the surface and charge complementarity between the two partners.

### ARL15 and CNNM co-localize in the perinuclear region and in kidney DCTs

The subcellular localization for the ARL15-CNNM complex was determined by immunocytochemistry in HEK293 cells. Overexpression of the mCherry-tagged Golgi-marker Beta-1,4-Galactosyltransferase 1 (B4GALT1) showed co-localization with ARL15 and CNNM2 in HEK293 cells, suggesting that the function of ARL15 is exerted within the perinuclear region containing the endoplasmic reticulum (ER) and the Golgi apparatus (Fig. 4A). Similar results were obtained with ARL15 and CNNM3, confirming that the proteins predominantly co-localize in the Golgi-apparatus (Fig. 4B). Since CNNM2 has been implicated in Mg^2+^ transport in the DCT segment of the kidney [7], we investigated whether ARL15 co-localizes with CNNM2 in this segment of the nephron (Fig. 4C). CNNM2 staining was concomitant with ARL15 staining in mouse kidney sections. In SK-RC-39 kidney cancer cells, similarly to HEK293, wild-type CNNM3 was co-localized with ARL15, however CNNM3 N-glycosylation mutant (N73A) did not co-localize with ARL15 (Fig. 4D). Wild-type CNNM3 and ARL15 co-localize partially with ER and Golgi and also showed co-localization at the plasma membrane (Fig. 4D). CNNM3 N73A mutant was mostly trapped in the ER and did not localize to plasma membrane (Fig. 4D). N-glycosylation of CNNM2 has been shown to regulate CNNM2 membrane trafficking [7] and it appears that it plays a similar role with CNNM3. One of the ARL15-interacting partners identified using BioID was ribophorin I protein (RPN1) (Supp. Fig. 1). RPN1 is part of the oligosaccharyltransferase complex that is involved in N-glycosylation and is found in the ER [19]. To confirm the interaction, co-immunoprecipitation assays with overexpressed CNNM3, ARL15 and RPN1 were performed (Supp. Fig. 4). CNNM3 interacted with RPN1 and ARL15, corroborating the localization of the three proteins to the ER as well (Fig. 4D). Our results demonstrate that ARL15 is localized in the ER and the Golgi-system and one of the enriched GO biological processes identified in ARL15 BioID was “Protein N-linked glycosylation via asparagine” (Fig. 1A), which suggests that ARL15 is involved in the N-glycosylation of CNNMs.

**Fig. 4.**
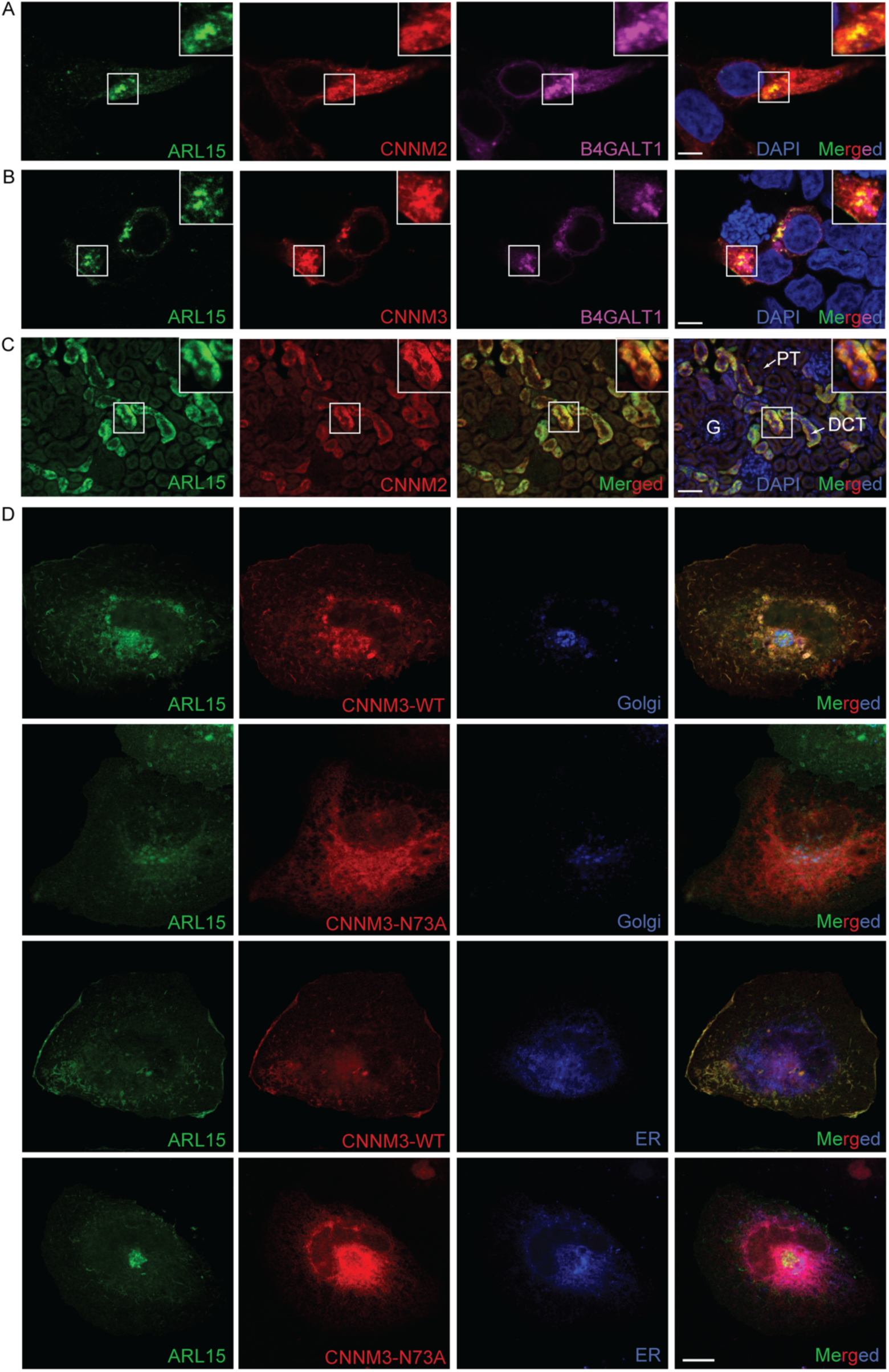
ARL15 and CNNMs co-localize in the Golgi system. **A**. Immunocytochemistry of HEK293 overexpressing ARL15, CNNM2 and mCherry-tagged Golgi-apparatus marker, B4GALT1. Nuclei are stained with DAPI. **B**. Immunocytochemistry of HEK293 overexpressing ARL15, CNNM3 and mCherry-tagged Golgi-apparatus marker, B4GALT1. Nuclei are stained with DAPI. **C**. Mouse kidneys were permeabilized and co-immunostained with anti-ARL15 and anti-CNNM2. Merged pictures stained for ARL15 in green, CNNM2 in red and DAPI (nuclei) in blue. Bar in figure A and B represents 5 μm, bar in figure C represents 50 μm. Representative image presented of three independent experiments with three replicates. G, Glomerulus, PT, Proximal tubule, DCT, Distal convoluted tubule. **D**. SK-RC-39 cells were co-transfected with ARL15-mClover3, CNNM3-mCherry (WT or N73A N-glycosylation mutant) and pmTurquoise2-ER or -Golgi

### CNNM3 N-glycosylation is modulated by ARL15 and Mg^2+^

As common post-translational protein modification, N-glycosylation plays important roles to regulate protein functions. Indeed, CNNM2 has been previously shown to have an N-glycosylation site close to its amino (N)-terminus, and it is necessary to stabilize CNNM2 on plasma membrane [7]. Using NetNGlyc [20], N-glycosylation sites were predicted in CNNM1, CNNM3 and CNNM4. The confirmed site in CNNM2 and the predicted sites in the three remaining CNNM family members were analogous (Supp. Fig. 5), all located in the extracellular N-terminal domain. To confirm that CNNM3 does indeed have an N-glycosylation site similar to CNNM2, the predicted asparagine 73 (N73) was mutated to alanine (N73A mutant). Compared to wild-type (WT) CNNM3, the N73A mutant migrated faster, which is indicative for the loss of N-glycosylation [21] (Fig. 5A). Endo H and PNGase F were used to characterize the type of glycans attached to CNNM3 (Fig. 5B). When WT CNNM3 was treated with Endo H, which cleaves high mannose and hybrid oligosaccharides from N-linked glycoproteins, the lower band of the doublet shifted further. Cleavage with PNGase F shifted all bands of CNNM3 to a lower molecular mass, indicating that the top band represents the hybrid/complex glycoform (Fig. 5A). To assess the effect of ARL15 on CNNM3 N-glycosylation, four different kidney cancer cell lines were used: Caki-1 and RCC4 as well as ACHN and SK-RC-39, belonging to renal clear cell carcinoma and papillary renal cell carcinoma, respectively. ARL15 was stably overexpressed in these four cell lines and the pattern of endogenous CNNM3 bands was evaluated. In all four cell lines, overexpression of ARL15 resulted in the increase of hybrid/complex glycoform and a decrease in oligomannose glycoform of CNNM3 (Fig. 5C). Interestingly, the effect of ARL15 expression on CNNM3 glycosylation could be modulated by the Mg^2+^ concentration in the growth media (Fig. 5D). When overexpressing ARL15 in SK-RC-39 cells, Mg^2+^ deficient medium resulted in an increase of the oligomannose CNNM3 glycoform. Additionally, CRISPR knockout of ARL15 resulted in a decrease of complex glycoform of CNNM3 as can be seen in the last two lanes (Fig. 5D).

**Fig. 5.**
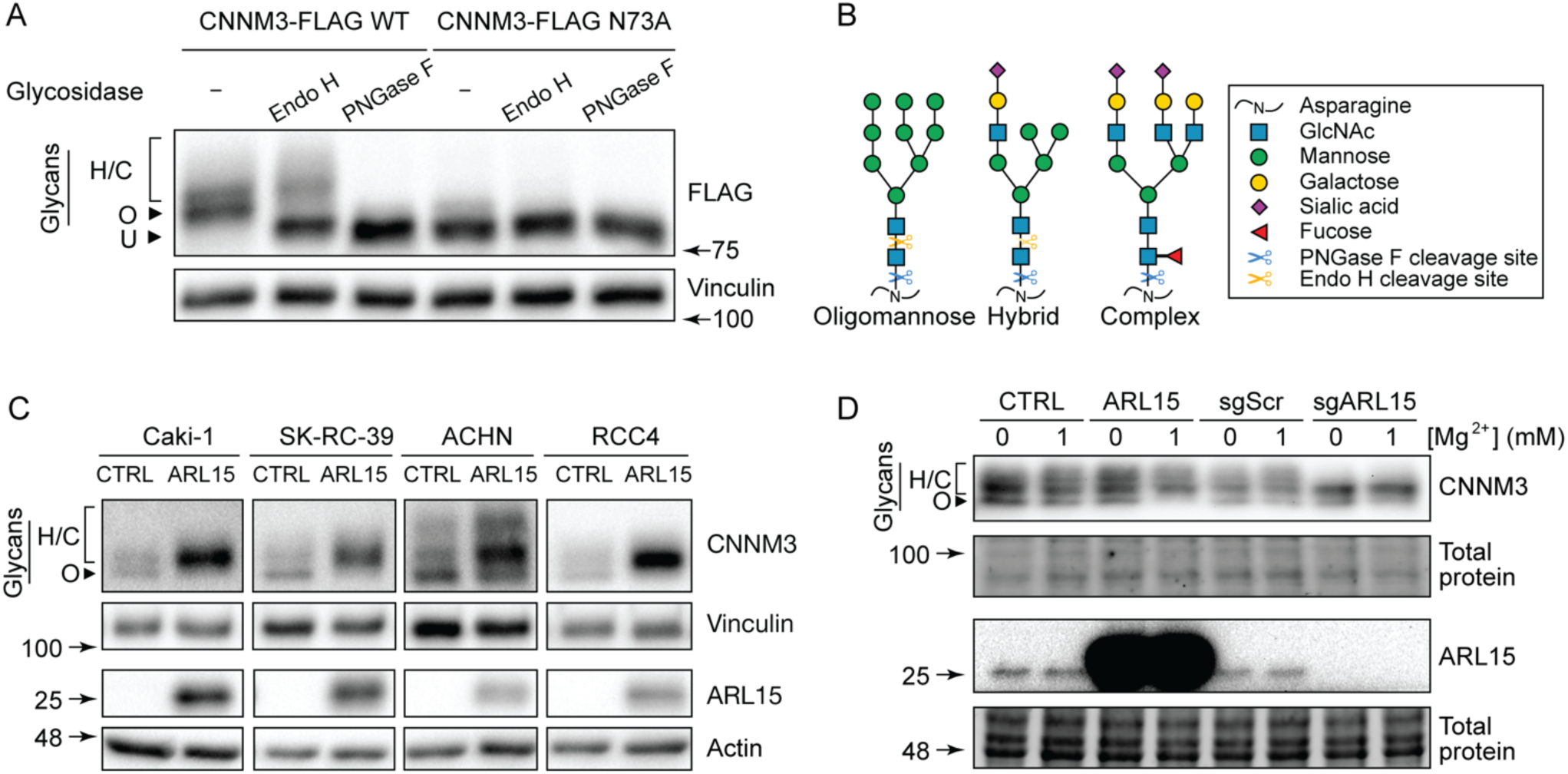
CNNM3 N-glycosylation is modulated by ARL15 and Mg^2+^ **A**. Wild-type and N-glycosylation mutant CNNM3 were treated with PNGase F and Endo H glycosidases to assess the presence of different glycoforms of CNNM3 and to confirm asparagine 73 as the site of glycosylation. **B**. Schematic representation of different types of glycans. **C**. Overexpression of ARL15 increases complex CNNM3 N-glycosylation in kidney cancer cells. **D**. SK-RC-39 cells were grown in the presence of absence of magnesium in the media and the status of CNNM3 glycoforms was assessed using western blotting CTRL, Control; sg, Guide RNA; Scr, Ccramble; ARL5, ADP ribosylation factor like GTPase 15; CNNM, Cyclin M; PNGase F, N-glycosidse F; Endo H, Endoglycosidase H.

### ARL15 affects Mg^2+^ flux and ATP production

To determine whether ARL15 regulates CNNM-dependent Mg^2+^ transport, ^25^Mg^2+^ uptake was studied. Kidney cancer cells were stably transduced to either overexpress or produce a CRISPR knockout of ARL15. ARL15 overexpression decreased ^25^Mg^2+^uptake, while ARL15 knockout significantly increased ^25^Mg^2+^ uptake more than 3-fold in RCC4 and SK-RC-39 cells (Fig. 6A). Since multiple enzymes involved in ATP production use Mg^2+^ as a co-factor and ATP is bound to intracellular Mg^2+^ to form Mg-ATP [1], we next assessed the impact of ARL15 expression on ATP production. In RCC4 and SK-RC-39 cells, overexpressing ARL15 resulted in a significant decrease in ATP production (Fig. 6B).

**Fig. 6.**
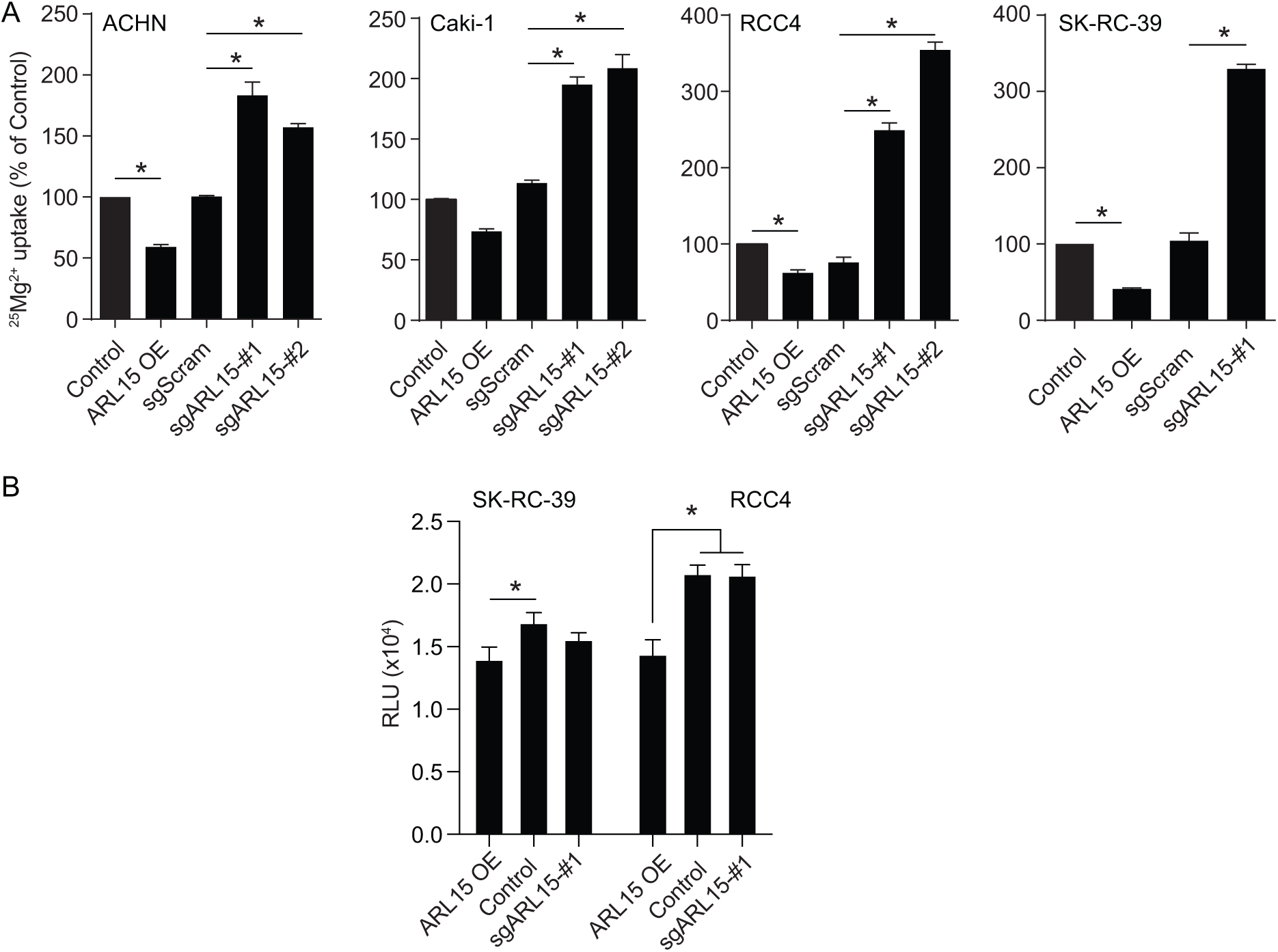
ARL15 affects Mg^2+^ flux and ATP production. **A**. ^25^Mg^2+^ uptake in stably overexpressed and knockdown ARL15 ACHN, Caki-1, RCC4, and SK-RC-39 cells. These various types of renal carcinoma cell lines were incubated with ^25^Mg^2+^ for 15 minutes and results were normalized to 0 minute. Each data represents the mean of 3 independent experiments ± SEM. * indicates significant difference compared to control cells. **B**. ATP production in stably overexpressed and knockdown ARL15 SK-RC-39 and RCC4 cells. Results are the mean ± SEM of 3 independent experiments. sg, Guide RNA; Scr, Scramble; OE, Overexpression; ARL5, ADP ribosylation factor like GTPase 15; CNNM, Cyclin M.

Endogenous cell surface expression of CNNM3 was further confirmed using cell surface biotinylation assay in SK-RC-39 cells with overexpression or CRISPR-mediated knockout of ARL15. The complex N-glycosylated CNNM3 glycoform migrates at approximately 90kDa, whereas oligomannose glycoform of CNNM3 migrates at around 75kDa. Although the plasma membrane expression of CNNM3 was not altered in the presence or absence of ARL15 (Supp. Fig. 6), mainly the oligomannose CNNM3 glycoform was observed at the plasma membrane in the ARL15 knockout cells.

## MATERIALS AND METHODS

### DNA constructs

Human *CNNM2* cDNA with an in-frame HA tag before CNNM2 stop codon and an *XhoI* restriction site after *CNNM2* stop codon was amplified using Phusion polymerase (New England Biolabs, Ipswich, MA, USA) and then the product was cloned into the pCINeo-IRES-GFP vector using *NheI* and *XhoI* (New England Biolabs). To obtain truncated *CNNM2* plasmids, primers (Forward : 5’-CGGCTAGCGCCACCATGATTGGCTGTGGCGCTTG-3’ and Reverse 1 (full-length *CNNM2*: 5’-CCCTCGAGCTATGCGTAGTCTGGCACGTCGTATGGGTAACCGGTGATGGCGCCTTCGTTG-3’), Reverse 2 (*CNNM2* transmembrane region: 5’-CCACCGGTCACGTCCTCCACCGTC-3’), Reverse 3 (*CNNM2* transmembrane+CBS1 region: 5’-CCACCGGTGTCATCGGGATCCAC-3’), Reverse 4 (*CNNM2* transmembrane+CBS1+link region: 5’-CCACCGGTGTGGTTATAAAATTTGGTGATG-3’) or Reverse 5 (*CNNM2* transmembrane+CBS1+link+CBS2 region: 5’-CGGCTAGCGCCACCATGATTGGCTGTGGCGCTTG-3’) were synthesized. All primers were purchased from Biolegio BV (Nijmegen, Netherlands). The PCR product was purified using a NucleoSpin® Gel and PCR Clean up kit (Macherey Nagel, Düren, Germany). Subsequently, the truncated *CNNM2* PCR products were cloned into the pCINE-IRES-GFP vector by digestion with restriction enzymes *NheI* (New England Biolabs) and *XhoI* (New England Biolabs).

To obtain C-terminal FLAG-tagged ARL15 and CNNM constructs, human coding sequences were amplified with appropriate primers containing attB1 and attB2 recombination site overhangs. Amplification was carried out with KAPA HiFi polymerase using GC buffer (Kapa Biosystems, Wilmington, MA, USA). Amplicons were gel purified and Gateway cloning was used to first recombine the amplicon into pDONR221 vector and then, pDEST26-C-FLAG (Addgene #79275) [22] vector, using Gateway BP or LR Clonase II enzyme mix, respectively (Thermo Fisher Scientific, Waltham, MA, US). Chemically competent DH5α *E. coli* were transformed with pDONR221 and pDEST26-C-FLAG vectors. To obtain C-terminal mClover3-tagged ARL15, mClover3 was amplified from pKanCMV-mClover3-18aa-actin vector (Addgene #74259) [23] and at the C-terminal of ARL15 vector using SLiCE cloning [24]. For fluorescence imaging of ER and Golgi pmTurquoise2-ER (Addgene #36204) and pmTurquoise2-Golgi (Addgene #36205) were used, respectively [25]. All constructs were verified by sequence analysis.

### Cell culture, transfection and transduction

HEK293 cells were grown in Dulbecco’s Modified Eagle Medium (DMEM) (Biowhittaker Europe, Vervier, Belgium) supplemented with 10% (v/v) fetal calf serum (PAA Laboratories, Linz, Austria), non-essential amino acids, and 2 mM L-glutamine at 37 °C in a humidified incubator with 5% (v/v) CO_2_ (New Brunswick Galaxy 170s). Cells were seeded 6 hours before transient transfection with Lipofectamine 2,000 (Invitrogen, Breda, The Netherlands), the ratio of DNA: Lipofectamine 2,000 was 1:2.

ACHN, Caki-1, HeLa, RCC4 and SK-RC-39 cells were grown in DMEM, high glucose (Thermo Fisher Scientific) supplemented with 10% fetal bovine serum (FBS) (Thermo Fisher Scientific) and 2 mM GlutaMAX at 37 °C in a humidified incubator with 5% (v/v) CO_2_.

For lentivirus production, HEK293T/17 cells were transfected with a 4:2:1 ratio of lentiviral construct of interest:PAX2:VSV-G. 24 hours later, the media was changed. 48 hours later, the media was collected and filtered through a 0.45 μm filter. To infect the cells, media was substituted with HEK293T/17 supernatant containing virus with 8 μg/ml polybrene. 48 hours later, the media was changed to fresh media containing a selection reagent and the selection was carried out for 7 days.

### Site-directed mutagenesis

To mutate asparagine 73 (N73) that was predicted to be the site of N-glycosylation of CNNM3 into alanine (A), pcDNA3.1-CNNM3-mCherry or pDONR221-CNNM3 carrying the wild-type sequence without a stop codon was amplified using a single primer PCR. Amplification was carried out with KAPA HiFi polymerase using GC buffer (Kapa Biosystems, Wilmington, MA, USA). The following primer was used: 5’-GGCCCGGGCTTCGCCgcCAGCTCTTGGTCCTGGGTGG-3’, indicating the nucleotides necessary to introduce the mutation in lower-case letters. After amplification, the original plasmid was digested with *DpnI* (New England Biolabs) for 2 hours at 37 °C, which was followed by transformation of chemically competent DH5α E. coli. The construct was verified by sequencing.

### Pull-down assay

A pull-down assay was performed to determine which proteins of mice DCT interact with CNNM2. pGEX-mCNNM2 c-tail-GST was inserted in bacteria to be able to multiply the protein. Afterwards bacteria were lysed with lysis buffer (150mM NaCl, 5mM EGTA, Triton 1% (v/v), 1mg/ml pepstatin, 1mM PMSF, 5mg/ml leupeptin, 5mg/ml aproptin, 50mM Tris/HCl, pH 7.5). Glutathione beads (Thermo Fisher, Rockford, IL, USA) were used to purify the samples to obtain only Cnnm2 c-tail. The beads were washed with pull down buffer (20mM Tris-HCL pH 7.4, 140mM NaCl, 1 mM CaCl_2_, 0.2% (v/v) triton-x-100, 0.2% (v/v) NP-40, 1:1000 pepstatin, 1:1000 Aprotin, 1:400 Leupeptin, 1:100 PMSF) and bacterial lysate was added to the beads followed by an incubation of 3 hours at 4 °C. After incubation, beads were transferred into a new tube and washed with pull down buffer. For kidney tissue, kidneys of 8 PV-GFP mice were dissected and minced in small pieces. The DCT was selected and dissolved in pull down buffer followed by homogenizing of the mixture using ultra turrax. After 30 minutes of incubation on ice, aliquots were taken and centrifuged. Supernatant was added to the beads followed by an incubation overnight rotating at 4 °C. After incubation, beads were washed with pull down buffer, Laemmli and DTT were added, and Western blot was performed. Migration was stopped when the sample was 1 cm in the running gel. Then samples were cut out and measured with mass spectrometry.

### BioID

BioID was performed as previously described [26]. Briefly, cleared lysates from ARL15^A86L^ expressing Flp-In™ T-REx™ HeLa cells were incubated with streptavidin beads (17-5113-01, 5 ml, GE Healthcare). Trypsin digestion on beads was performed and peptides were collected in water, dried in a SpeedVac and pellets were resuspended in 5% formic acid prior to their injection into the mass spectrometers. Digested peptides from HEK293 and HeLa cells were respectively injected into LTQ-Orbitrap Velos and Q Exactive (Thermo Fisher) onto a 75 μm i.d. × 150 mm Self-Pack C18 column installed in the Easy-nLC 1000 system (Proxeon Biosystems) coupled to a Nanospray Flex Ion Source, at the IRCM Proteomics core facility.

Raw mass spectrometry files were analyzed, as previously described [26], using the Human RefSeq database (version 57) supplemented with ‘‘common contaminants’’ from Max Planck Institute (https://maxquant.org/), Global Proteome Machine (http://www.thegpm.org/crap/index.html) and decoy sequences with the Mascot search engine through the iProphet pipeline integrated in ProHits. Significance Analysis of INTeractome (SAINT; version 3.6.1) files generated with ProHits using iProphet protein probability ≥ 0.9 and unique peptides ≥ 2, comparing samples against negative controls, were used to estimate interactions statistics with ProHits-viz [27]. (https://prohits-viz.lunenfeld.ca/). An average probability (AvgP) ≥ 0.95 was used as a cutoff to consider statistically significant and of high confidence interactions. Gene ontology overrepresentation analysis was performed using PANTHER 15.0 [28].

### Gel filtration

Gel filtration chromatography of the CNNM2_BAT_.ARL15 complex was performed using 1 mg/mL protein at 0.5 ml/min using a Superdex 200 Increase 10/300 GL column in 50 mM Hepes, pH 8.5, 200 mM NaCl, 1 mM MgCl_2_, 1 mM TCEP buffer. The following values were used: V_o_=8.61 mL, void volume; V_t_=24 mL, total column volume.The calibration trendline is *y*= −0.2451*x* + 1.5726 (*y* = *K*_av_; *x* = log*M*). The standard proteins used to calibrate the column were: 1, thyroglobulin; 2, gamma-globulin; 3, ovoalbumin; 4, myoglobin; 6, Vitamin B12.

### ARL15 model

A model of the structure of ARL15 was generated by simulated annealing as implemented in MODELLER [29] using as templates the X-ray diffraction coordinates of the complex between human ARL2 and BART (pdb code 3DOE chain A; 36.3 identity) complex between murine ARL2 and PDEδ (pdb code 1KSG chain A; 36.3 identity) murine ARF6 (pdb code 6BBQ; 35.67% identity) human ARF6 (pdb code 2A5G; 35.67% identity). Results were monitored in UCSF Chimera [30]. A single structure—displaying the lowest zDOPE [31] score (−0.58) and an RMSD value of 3.54 Å with respect to the closest template—out of 100 results was selected. This structure was subjected energy minimization using the Amber 14SB force field [32]. A final check was performed with Coot [33] before deposition and the ModelArchive server.

### CNNM2 models

To generate alternative linker conformations, in addition to completing sidechains from a SAXS model, simulated annealing computations were carried out using MODELLER [29]. 100 conformations were generated, arranged according to their zDOPE [31] scores and tested for collisions with other subunits with UCSF Chimera [30]. At the end, three different conformations were tested. A model of the cytosolic domains of CNNM3 was also generated using the full precursor sequence (NCBI accession code 060093.3) and the XRD coordinates of the CNNM2 and CNNM4 cytosolic domains, obtained from corresponding crystals of the independent cytosolic domains and from SAXS data of the entire cytosolic region of CNNM4 [34]. The dimeric assembly of the complementary Bateman modules used in the model corresponds to the flat (ATP-bound and/or PRL-bound like) conformation of the CBS module (association of two Bateman domains) [34][35]. The modeled region showed a 53.6 % sequence identity with respect to the template. Best model showed 6.1 Á RMSD with respect to the template, and a zDOPE value of 0.39. This positive value is mainly due to the model including flexible loop and linker regions.

### Docking models

Docking computations were carried out with benchtop HEX [36] and compared with Brownian dynamics (BD) computations (see Supplemental methods). Hex computations included shape evaluation, in vacuo electrostatics and Decoys As a Reference State (DARS) potentials [37]. In every simulation, 20000 structures were generated, clustered using a RMSD cut-off value of 3 Å and arranged according to their energies. First 100 were selected for analysis. Post-processing consisted in a short OPLS energy minimization as implemented in the software.

### Brownian dynamics

Docking modeling was complemented with Brownian Dynamics computations using the WebSDA server [38] to test if results could be reproduced by other methods. The force-field grids used in BD included all electrostatics and desolvation [39][40] grids. Charges were added with PDB2PQR [41].

200 trajectories of BD computations were run in docking mode. A total of 500 non-redundant complexes were recorded along the computations. Supplemental table YD displays the summary of the clustering analysis of the distinct complexes found. First cluster corresponds to binding to CNBH domains at regions nearby missing structure, so they are considered artefactual. Cluster 2, however, showed the highest population and—on average— displayed lowest energies, and the lowest dispersion of conformations, according to RMSD values with respect to the representative structure. Supplemental figure ZD illustrates the consistency between the BD computations and docking simulations performed with Hex (see main text).

### Glycosidase and tunicamycin treatment

20 μg of protein from SK-RC-39 cells were treated with N-glycosidase F (PNGase F) and Endoglycosidase H (Endo H) (New England Biolabs) according to the manufacturer’s instructions. Briefly: protein was denatured at 100 °C for 10 minutes, 1 unit/μl of a glycosidase was added with appropriate buffers and digested at 37 °C for 1 hour. SK-RC-39 cells grown in DMEM were treated with tunicamycin (Sigma-Aldrich) at 1 ug/ml for 8 hours to inhibit N-glycosylation.

### Immunohistochemistry

5 μm kidney frozen sections were fixed with formalin and washed with TN buffer (0.1 M Tris/HCl (pH 7.6) 0.15 M NaCl). Then, the sections were permeabilized in TN-Triton (TN with 0.1% (v/v) Triton X-100) for 30 minutes. After incubation, the sections were washed and blocked with TN with 0.5% (w/v) Blocking Reagent TSA fluorescein system (Perkin Elmer, Waltham, MA, USA) for 30 minutes. The sections were incubated overnight at 4 °C with the following primary antibodies: guinea pig anti-CNNM2 (1:100), rabbit anti-ARL15 (Sigma-Aldrich, St. Louis, MO, USA, 1:50). For detection, kidney sections were incubated with Alexa Fluor 488 conjugated goat anti-rabbit and Alexa Fluor 596 conjugated goat anti-Guinea pig IgG secondary antibodies (Thermo Fisher, Rockford, IL, USA, 1:300). Images were visualized using AxioCam cameras (Zeiss, Oberkochen, Germany) and AxioVision software (Zeiss).

### Immunocytochemistry

HEK293 cells were seeded at a low density on 0.01% (w/v) poly-L-lysine (Sigma, St Louis, MO, USA) coated coverslips and were transfected the subsequent day using Lipofectamine 2000 (Invitrogen, Breda, The Netherlands). Cells were transfected with the following constructs: pLenti6-CNNM3-V5, pDEST26-ARL15-FLAG, and mCherry-Golgi-7. After 24 hours, cells were washed with PBS, followed by fixation for 10 minutes using 4% (w/v) methanol-free formaldehyde (ThermoFisher, Waltham, MA, USA) in PBS. Afterwards, cells were permeabilized for 10 minutes in 0.1% (v/v) Triton-X100 and 0.3% (w/v) bovine serum albumin solution. Subsequently, cells were treated with 50 mM NH_4_Cl in PBS to reduce background. Thereafter, cells were washed 3x in PBS followed by blocking in 16% (v/v) normal goat serum (Merck Milipore, Burlington, MA, USA) supplemented with 0.1% (v/v) Triton-X100 for 30 minutes. Cells were then probed with primary antibody diluted in blocking buffer overnight at 4 °C. Cells were washed three times and incubated in secondary antibodies for 45 minutes at room temperature in the dark. Cells were washed three times with PBS and subsequently mounted using mounting medium supplemented with 4′, 6-diamidino-2-phenylindole (DAPI; Southern Biotech, Birmingham, AL, USA). Images were made using the Zeiss LSM880 (Oberkochen, Germany) and analysed using the freely available Fiji software [42]. The following antibodies were used: mouse monoclonal anti-V5 (Thermo Fisher, Rockford, IL, USA, 1:1,000) and rabbit polyclonal anti-FLAG (Sigma-Aldrich, St. Louis, MO, USA, 1:1,000), Alexa Fluor 488 conjugated goat anti-rabbit and Alexa Fluor 647 conjugated goat anti-mouse (Thermo Fisher, Rockford, IL, USA, 1:300).

### Fluorescence imaging

SK-RC-39 cells were transfected with pcDNA3.1-CNNM3-WT-mCherry WT, pcDNA3.1-CNNM3-N73A-mCherry, ARL15-mClover3, pmTurquoise2-ER and pmTurquoise2-Golgi using Lipofectamine 2000 (Invitrogen, Breda, The Netherlands) for 24H and then transferred to an 8 chamber slide (ibidi GmbH, Fitchburg, WI, USA). Cells were fixed with 4%PFA for 10 min, briefly rinsed with PBS, then coverslips were mounted with Mowiol mounting medium (Millipore Sigma, Darmstadt, Germany). Slides were cured for 24 hrs and pictures were taken by using the Zeiss LSM880 with Airyscan processing (Zeiss, Oberkochen, Germany).

### Western blot

Cells were lysed with lysis buffer added 1 mg/ml pepstatin, 1 mM PMSF, 5 mg/ml leupeptin, 5 mg/ml aproptin protease inhibitors. The lysis buffer contained 50 mM Tris-HCl (pH 7.5), 1 mM EDTA, 1 mM EGTA, 150 mM NaCl, 1 mM sodium-orthovanadate, 10 mM sodium-glycerophosphate, 50 mM sodium fluoride, 0.27 M sucrose, 10 mM sodium pyrophosphate, 1% (v/v) Triton X-100. Protein concentrations were measured by performing a Bicinchoninic Acid protein assay (BCA) (Fisher Scientific, Hampton, NH, USA) and 20 µg protein was used for western blot. Blots were then incubated in primary antibody overnight rolling at 4 °C. Primary antibodies were diluted in 1% (w/v) milk diluted in TBST as following; anti-HA (Cell Signaling technology, Leiden, The Netherlands) diluted 1:5,000, anti-FLAG (Sigma-Aldrich, St. Louis, MO, USA) diluted 1:5,000, anti-V5 (Thermo Fisher, Rockford, IL, USA) diluted 1:5,000, anti-β-actin (Sigma-Aldrich, St. Louis, MO, USA) diluted 1:10,000, all are raised in mice. After washing with Tris-Buffered Saline and Tween 20 (TBST), blots were incubated for 1 hour rolling at RT in secondary antibody which was anti-mouse antibody raised in sheep (Sigma-Aldrich, St. Louis, MO, USA) diluted 1:10,000. Then, proteins were visualized with were made with ChemiDoc (Bio-Rad Laboratories Inc, Hercules, CA, USA). Subsequent analysis was done using ImageJ [42].

### Co-Immunoprecipitation

HEK293 or HeLa cells were seeded and transfected in petri dishes, 24 to 48 hours after transfection cells were lysed with lysis buffer. 12 hours before collecting the transfected HEK293 cells, carbobenzoxy-Leu-Leu-leucinal (MG132) (Sigma-Aldrich, St. Louis, MO, USA) dissolved in Dimethyl sulfoxide (DMSO) were treated to the HEK293 cells. BCA assay was performed to determine the protein concentration of the lysates. Input samples were taken to be able to check transfection efficiency by performing western blot. For co-immunoprecipitation samples, 30 µl/sample protein A/G plus agarose beads (Santa Cruz Biotechnology, CA, USA) were washed with PBS. And 1.5 µl/sample in mouse raised antibody for human HA (Sigma-Aldrich, St. Louis, MO, USA) was added to the beads. For negative control, no antibody was added to the beads. Samples were then incubated for 2 hours rotated at 4 °C. Unbound antibodies were removed by washing with lysis buffer. Then lysates with the same amount of proteins (as calculated with the BCA test) were added to the beads and incubated overnight rotated at 4 °C. After incubation, lysates and beads were washed with PBS, Laemmli and DTT were added, and western blot was performed. For FLAG-tagged protein immunoprecipitation, for each sample 20 μg of ANTI-FLAG® M2 affinity agarose gel (Sigma-Aldrich, St. Louis, MO, US) were pre-blocked with 5% (w/v) BSA and 300 ug of total cell lysate (TCL) was pre-cleared with 20 μg of Protein A/G agarose mix, for 2H at 4°C on a rotator. TCL supernatant was collected and incubated with pre-blocked FLAG beads for 2 hours at 4°C on a rotator.

### CRISPR-Cas9 experiments

sgRNAs targeting ARL15 were cloned into lentiCRISPRv2 plasmid (Addgene #52961) [43] using Golden-Gate Assembly with the Esp3I endonuclease following the standard protocol [44]. For lentivirus production, HEK293T/17 were transfected using Lipofectamine 2000 (Invitrogen, Breda, The Netherlands) with 4:2:1 ratio of lentiCRISPRv2:PAX2:VSV-G. 24H later the media was changed. 48H later the media was collected and filtered through a 0.45 μm filter. To transduce the cells, media was substituted with HEK293T/17 supernatant containing virus with 8 μg/ml polybrene. 48H later the media was changed to fresh media containing puromycin and the selection was carried out for 7 days.

### ATP production

5,000 kidney cancer cells were plated per well in a 96-well plate and ATP production was measured after growing overnight. CellTiter-Glo 2 kit (Promega) using manufacturer’s instructions.

### ^25^Mg^2+^ uptake

Cells were seeded in petri dishes and 24 hours after transfection cells were reseeded into 12-well plates. Two days after transfection, cells were washed once with basic uptake buffer without Mg^2+^ (125 mM NaCl, 5 mM KCl, 0.5 mM CaCl_2_, 0.5 mM Na_2_HPO_4_, 0.5 mM Na_2_SO_4_, 15 mM HEPES/NaOH, pH 7.5), followed by an incubation for 0 or 15 minutes at 37°C with basic uptake buffer containing 1 mM ^25^Mg^2+^ (Cortecnet, Voisins Le Bretonneux, France). After incubation, the basic uptake buffer containing ^25^Mg^2+^ was removed and cells were washed with ice cold PBS. Then cells were lysed with 1 ml nitric acid which was then diluted in Milli-Q until a nitric acid (Sigma Aldrich, St. Louis, MO, USA) concentration of 10%. Samples were analysed with ICP-MS (inductively coupled plasma mass spectrometry).

### Cell surface biotinylation

Petri dishes were coated with poly-L-lysine before seeding and transfection. Two days after transfection the cells were washed with PBS-CM (100 ml 10x PBS, 1 mM MgCl_2_, 0.5 mM CaCl_2_, pH 8.0 adjusted with NaOH, Milli-Q until total volume of 1,000 ml) followed by adding 0.5 mg/ml Sulfo-NHS-LC-LC-biotin (Thermo fisher scientific, Rockford, USA). After 30 minutes incubation, cells were washed with 0.1% (w/v) PBS-CM BSA and PBS. Cells were then lysed with lysis buffer, and protein concentration was measured with a BCA test. Input samples were taken to be able to check transfection efficiency by performing Western blot. Then, lysates with the same amount of proteins (as calculated with the BCA test) were added with the neutravidin agarose beads (Thermo fisher scientific, Rockford, USA) and incubated overnight rotated at 4°C. After incubation, protein lysates were washed with lysis buffer and Laemmli and DTT buffer were added and incubated at 37°C for 30 minutes before loading the protein samples on SDS-PAGE.

### Statistical analysis

Results are expressed as mean ± standard error of the mean (SEM). Biotinylation results of experiments are statistically analyzed by performing one-way ANOVA followed by Tukey as post-test. ^25^Mg^2+^ uptake results with CNNM2 and ARL15 are statistically analyzed by performing a two-way ANOVA followed by Tukey as post-test. Differences with p<0.05 were regarded as statistically significant.

## DISCUSSION

In this study, we identified ARL15 as a novel interacting partner of CNNMs. ARL15 binds to CNNMs in the ER and regulates their complex N-glycosylation in the Golgi system (Fig. 7). Our data suggest that the complex N-glycosylation of CNNMs reduces their activity at the plasma membrane, as reflected in a reduced ^25^Mg^2+^ uptake in renal carcinoma cells. As such, ARL15 may affect cellular Mg^2+^ homeostasis and energy metabolism.

**Fig. 7.**
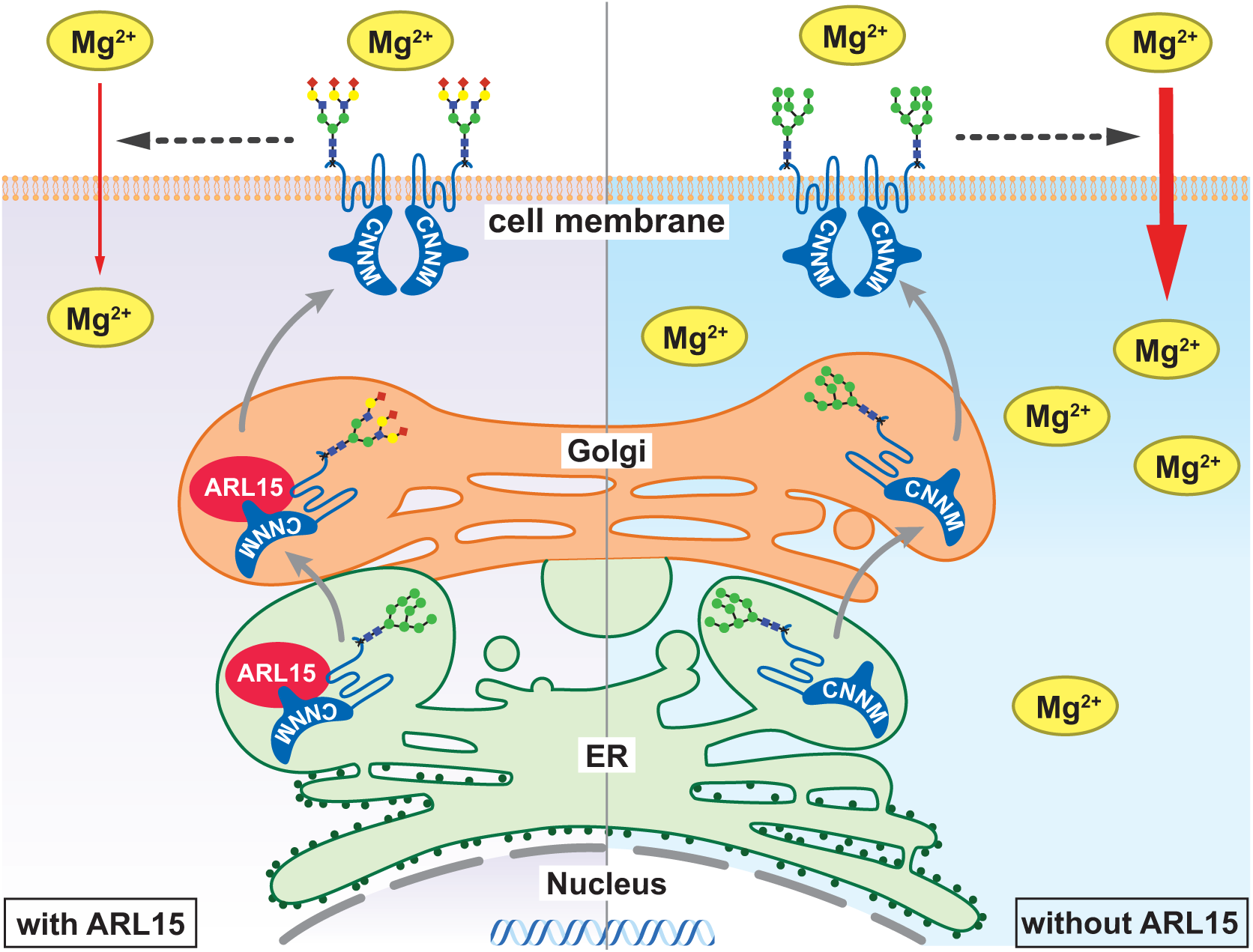
Summary of the effect of ARL15•CNNM complex on Mg^2+^ flux. In the presence of ARL15, it interacts with CNNMs in the ER and Golgi, resulting in their complex N-glycosylation, which in turn decreases Mg^2+^ uptake. In the absence of ARL15, CNNMs are found in less complex glycoforms and that results in increased Mg^2+^ uptake. ER, Endoplasmic reticulum; ARL5, ADP ribosylation factor like GTPase 15; CNNM, Cyclin M.

ARL15 is a member of the superfamily of ARF-like (ARL) proteins, which have been functionally characterized as small GTPases. Although ARL15 has been relatively scarcely studied to date, many studies have demonstrated that other ARL and ARF proteins are involved in vesicle trafficking [45]. Our immunocytochemistry in HEK293 cells demonstrated that ARL15 is predominantly localized in the Golgi system (Fig. 4), which is in line with previous reports of ARL15 in 3T3-L1 pre-adipocytes [46]. Moreover, proximity labeling of ARL15 identified many Golgi-specific proteins (Supp. Fig. 1). However, the localization with RPN1, a protein of the oligosaccharyltransferase complex, suggests that the interaction between ARL15 and CNNMs is already formed in the ER. Indeed, several ARF proteins, e.g. ARF1 and ARF3, have been shown to regulate ER-to-Golgi trafficking [47].

The main finding of our study is that ARL15 binds CNNMs and modulates their activity via N-glycosylation. Biosynthesis of glycoproteins commences in the ER, where a pre-assembled oligosaccharide is bound onto the nascent protein by oligosaccharyltransferases. After passing the ER, a series of reactions further assemble the glycan in the medial-Golgi and maturation takes place in the trans-Golgi. Three different types of N-glycans are commonly present on proteins; oligomannose, hybrid and complex N-glycans. Our results demonstrate that ARL15 stimulates the formation of complex N-glycans on CNNMs. Although ARL15 does not possess the enzymatic capacity to change the glycosylation *per se*, we have identified several proteins of the oligosaccharyltransferase complex in our BioID experiments (e.g. RPN1, STT3A and STT3B) (Fig. 1). Moreover, protein N-glycosylation was among the enriched GO-terms in our analysis. We hypothesize that trans-Golgi trafficking via ARL15 is an essential step in the complex glycosylation of CNNMs. A similar mechanism has been postulated for ARFGEF1, which regulates the N-glycosylation and trafficking of integrins [48].

The N-glycosylation of CNNMs is essential for their plasma membrane expression. CNNMs contain a single N-glycosylation site in the extracellular N-terminal region [7]. Mutation of the Asn73 residue to Ala in CNNM3 completely abrogated its membrane expression (Fig. 5A). Similar results were previously obtained for Asn112 residue in CNNM2 [7]. In this study, we demonstrated that not only the presence of, but also the composition of the glycan is essential for the protein function. ARL15 overexpression resulted in a complex glycosylation of CNNM3, which was accompanied by a significant reduction of ^25^Mg^2+^ uptake. *Vice versa*, ARL15 downregulation resulted in oligomannose glycoform and increased ^25^Mg^2+^ uptake. Altogether, we propose a model in which the composition of the N-glycan affects CNNM activity (Summarized in Figure 7).

Multiple roles for N-glycosylation of plasma membrane proteins have been described, ranging from effects on membrane trafficking, membrane stability and protein degradation [49][50]. In our experiments, neither ARL15 overexpression nor ARL15 downregulation affected the plasma membrane localization of CNNM3 (Supp. Fig. 4), suggesting that membrane trafficking of CNNM3 is not affected by ARL15. Consequently, changes in N-glycosylation may directly affect the activity of CNNM3 at the plasma membrane. It has been extensively described that N-glycans interact with glycan-binding proteins in the extracellular spaces, such as lectins. Lectin-glycan interactions are involved in many biological cellular processes such as apoptosis, differentiation as well as regulation of membrane transport [51]. Indeed, binding of lectins and other glycan-binding proteins has been shown to regulate TRPV5-mediated Ca^2+^ transport [52][53]. Whether the binding of lectins explains the changes in CNNM3 activity remains to be defined.

The interaction model that we developed is compatible with a 1:1 CNNM2:ARL15 stoichiometry. ARL15 interacts with the Bateman CBS1 domains, in particular with a surface region defined between α−helix A, the β1-β2 hairpin loop and α-helix H1. Indeed, co-immunoprecipitation experiments confirmed that the CBS domains are essential for ARL15 binding (Fig. 2B). The interaction also involves the CNBH domains 3C, thereby explaining the larger affinity observed with the full-length constructs. Overall, the interaction displays high surface complementarity and presents a core of hydrophobic and polar residues surrounded by charged ones. Interestingly, the negative-charged residues present in the CNNM2 docking site is complementarity to the positive-charged surface of ARL15 (Fig. 3D). The interaction of ARL15 with CNNM3 cytosolic domain seems to use the same region. The more negative charge as compared to CNNM2 may explain the larger affinity toward positively charged ARL15. Notably, no overlap is observed between the binding site of ARL15 and PRL, which mostly binds CBS2 by its long loop region comprising β5 and β6 [35] (Supp. Movie 1).

The identification of ARL15 as a novel regulator of CNNM Mg^2+^ transport activity is of particular interest in the DCT segment of the kidney. In the DCT, transient receptor potential cation channel subfamily M member 6 (TRPM6) facilitates apical Mg^2+^ transport [54]. Basolateral Mg^2+^ extrusion in the DCT is regulated by CNNM2. Consequently, mutations in CNNM2 and TRPM6 have been shown to cause renal Mg^2+^ wasting in patients [8][55][56]. Interestingly, the ARL15 locus was recently associated with urinary Mg^2+^ wasting in a GWAS [15]. In the same study, TRPM6 channel activity was shown to be significantly increased in the presence of ARL15 [15]. Our results demonstrate that ARL15 also binds CNNM2, suggesting that both the apical and basolateral Mg^2+^ transport mechanisms are simultaneously regulated. These findings explain how ARL15 determines urinary Mg^2+^ excretion.

Interestingly, overexpression of ARL15 resulted in decreased ATP levels (Fig. 6B). Intracellular Mg^2+^ and ATP levels are closely associated, as ATP must be bound to this cation to be biologically active [1]. Indeed, previous studies have shown that PRL-2 knockdown and decreased intracellular Mg^2+^ levels, reduce intracellular ATP levels and regulate cellular metabolism [3][57]. Given that ARL15 overexpression reduced CNNM-mediated Mg^2+^ uptake, the expression of ARL15 may indirectly regulate cellular metabolism. Indeed, ARL15 has been associated with a wide range of metabolic parameters and diseases in GWAS, including adiponectin, HDL, diabetes mellitus and body shape [58][59][60][61][62][63]. As these studies did not analyze Mg^2+^ status as a modifying factor, it cannot be excluded that ARL15 has additional functions that explain these associations. Indeed, ARL15 has been demonstrated to modify the insulin-signaling pathway in myotubes [64].

Overall, our work establishes complex N-glycosylation of CNNMs as an essential process to regulate their activity. This crucial post-translational modification promoted by ARL15 on CNNMs adds to recent mechanisms of CNNMs modulation such as their circadian rhythm expression [65] and their interactions with the PRL family by a magnesium-sensitive mechanism [3][11]. The increasing complexity of CNNMs regulation provides a dynamic system to ensure the correct levels of intracellular Mg^2+^ during metabolic changes and cell requirements.

## DATA AVAILABILITY

ARL15 model is available at ModelArchive server DOI: 10.5452/ma-c4ded. CNNM3 model is available at ModelArchive server DOI 10.5452/ma-v6la8 The BioID data will deposited to ProteomeXchange upon acceptance of the manuscript.

## ACKNOWLEDGEMENTS

We thank Caro Bos, Milou Smits and Shirin Mostert for excellent technical support. This work was financially supported by grants from the European Joint Program for Rare Diseases (EJPRD2019-40), the Netherlands Organization for Scientific Research (NWO Veni 016.186.012. Vici 016.130.668), the Dutch Diabetes Research Foundation (2017.81.014), the Spanish Ministry of Science and Innovation and Universities (PGC2018-096049-B-I00), European Regional Development Fund (FEDER), and the Andalusian Government (BIO-198, US-1254317 and US-1257019). This project “FIGHT-CNNM2” was supported by a grant from Fonds de recherche du Québec -Santé (FRQS) FRQ-S #294027 and by a grant from the CIHR ERT-168503, under the frame of E-Rare-3, the ERA-Net for Research on Rare Diseases. In addition, this project has received funding from the European Union’s Horizon 2020 research and innovation programme under the EJP RD COFUND-EJP N° 825575.

In addition, it was also supported by Canadian Institutes of Health Research Grants MOP-142497 and FDN-159923. M.L.T. is a Jeanne and Jean-Louis Lévesque Chair in Cancer Research and holds a Distinguished James McGill Chair from McGill University. Y.Z. was supported by a scholarship from the Fonds de recherche du Québec – Santé (FRQS) and by a travel award from the Canderel Foundation at the Goodman Cancer Research Centre.

## AUTHOR CONTRIBUTIONS

Y.Z., C.M., S.H., J.H., M.L.T. and J.d.B. designed the research studies, Y.Z., C.M., I.D.M., A.D.Q., G.F., I.G.R., E.K., F.L., J.B., M.-P.T., L.A.M.C., J.F.C., N.U., S.H., J.H., M.L.T. and J.d.B. conducted experiments and/or analyzed data, Y.Z., C.M., S.H., M.L.T. and J.d.B. wrote the manuscript. All authors corrected the manuscript and approved the final version.

