## Supplemental Figures and Tables for "ARL15 modulates magnesium homeostasis through N-glycosylation of CNNMs"

### Supplementary Figures

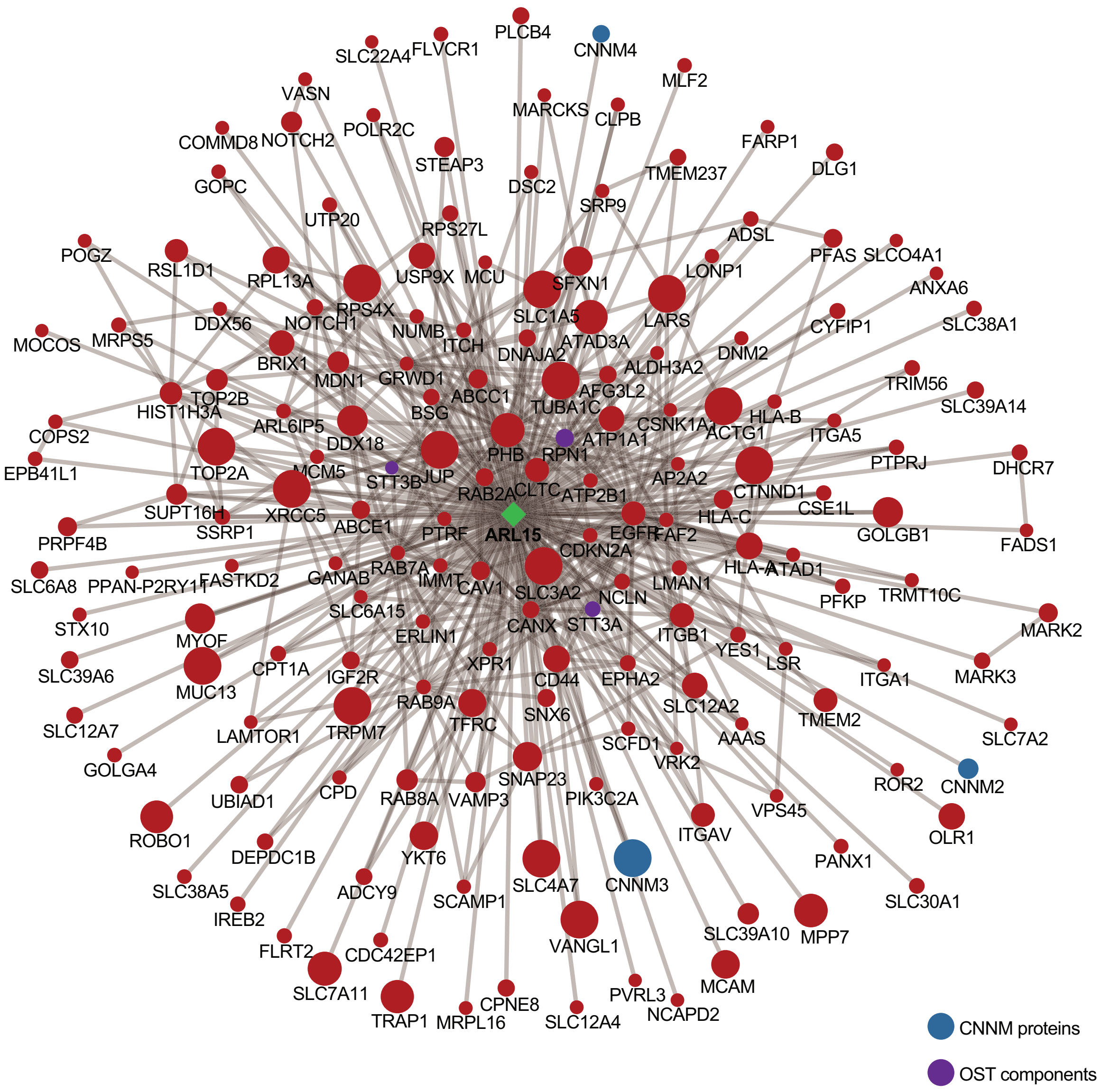


**Supplementary Figure 1: ARL15 interacting partners identified through BioID**

ARL15 interacting partners are presented with the size of the circle corresponding to the average spectral count and the edges between ARL15 and preys were obtained by augmenting the network using 3 different databases; BioGRID, IntAct and iRefIndex.

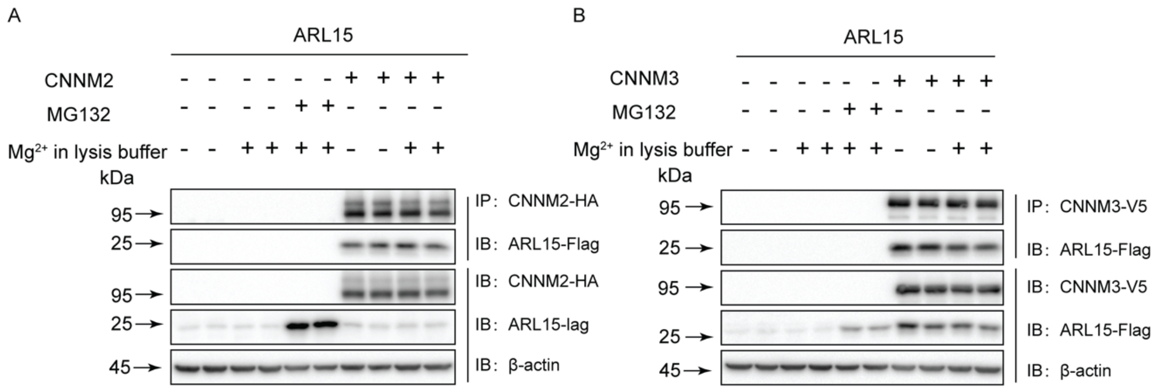


**Supp. Figure 2. Binding of ARL15 and CNNM2/3 is independent of magnesium**

**A.** Co-immunoprecipitation of HEK293 cells transfected with ARL15-FLAG and HA-CNNM2 treated with or without 1mM $Mg^{2+}$ in lysis buffer. The upper two blots show the detection of the FLAG-tagged proteins in anti-HA precipitated cell lysates. The lower two blots show input controls of HA-tagged and Flag-tagged proteins respectively.

**B.** Co-immunoprecipitation of HEK293 cells transfected with ARL15-FLAG and V5-CNNM3 treated with or without 1mM $Mg^{2+}$ in lysis buffer. The upper two blots show the detection of the FLAG-tagged proteins in anti-V5 precipitated cell lysates. The lower two blots show input controls of V5-tagged and FLAG-tagged proteins respectively. The figure shows a representative blot of 3 independent experiments.

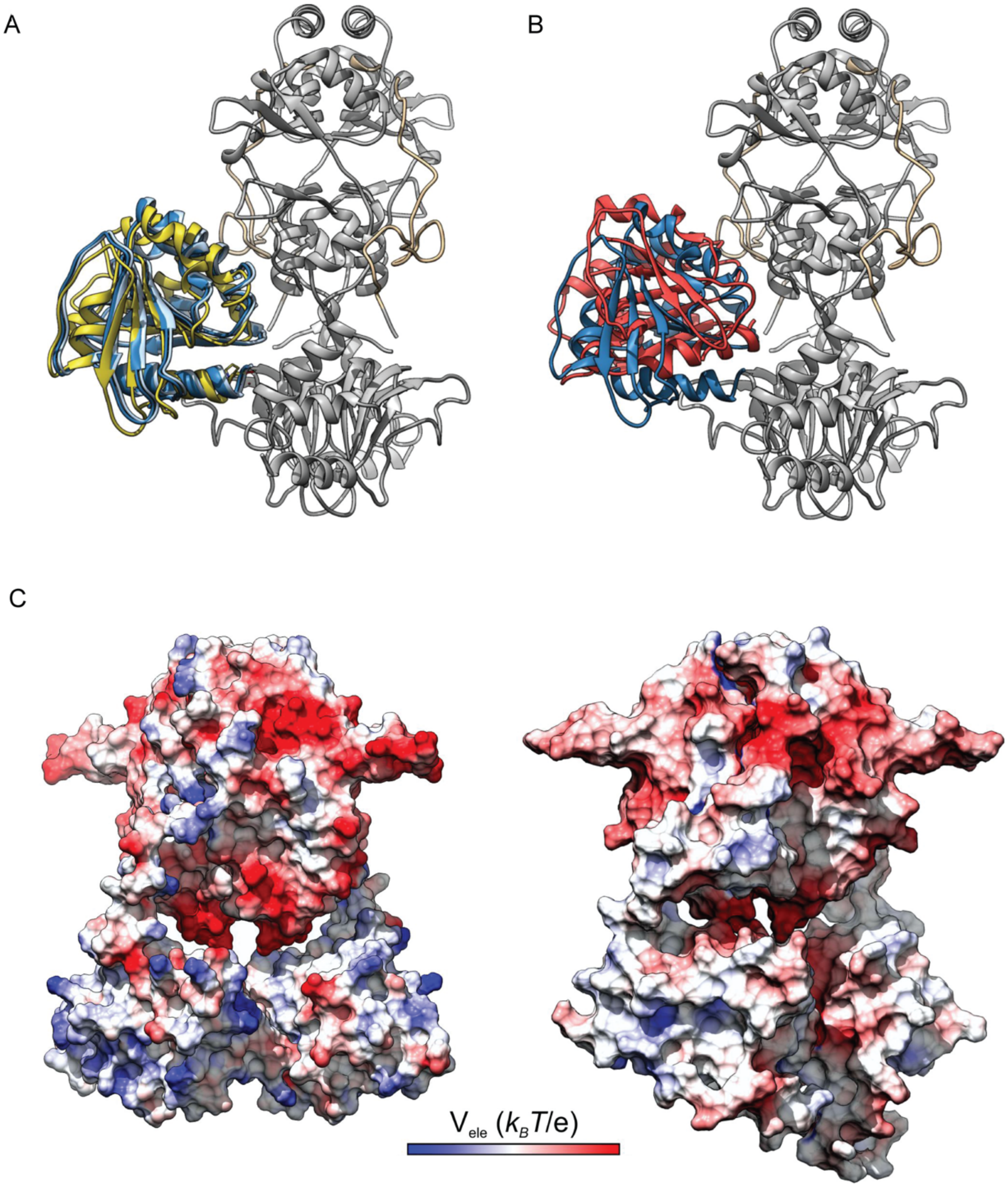


**Supp. Figure 3 Overlay of different CNNM2 and CNNM3 docking models**

**A.** Overlay of two solutions (dark and light cyan) from different HEX docking computations with the representative structure of the most populated cluster (yellow) from Brownian dynamics computations.

**B.** Overlay of the results from HEX docking computations between Arl15 and CNNM2 and

CNNM3. CNNM3 was aligned to CNNM2—which is shown. The ARL15 bound to CNNM3 is in red ribbons.

**C.** Comparison of CNNM2 and CNNM3 electrostatic potentials at the surface level. Positive potentials are in blue, and negative in red. Color scales as in Fig. 3B. The CNNM3 model includes long disordered loops in the CNBH domain loops at the bottom of the structure.

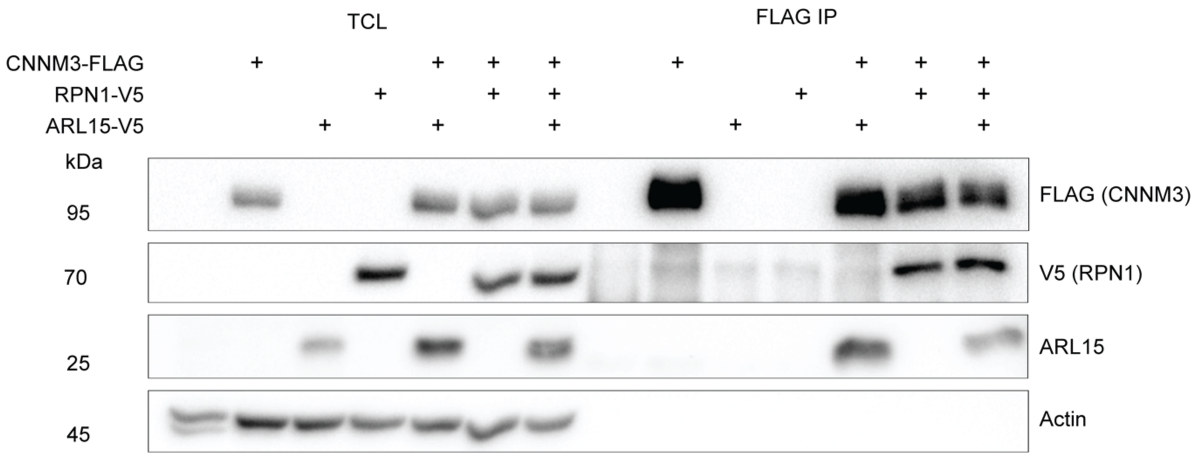


**Supp. Figure 4 Co-immunoprecipitation of CNNM3, ARL15 and RPN1**

HeLa cells were co-transfected with CNMM3-FLAG, ARL15 and RPN1-V5 and immunoprecipitation was performed using magnetic FLAG beads. CNNM3 co-immunoprecipitated with ARL15 and RPN1.

TCL, Total cell lysate; IP, Immunoprecipitation; ARL15, ADP ribosylation factor like GTPase 15; CNNM, Cyclin M; RPN1, Ribophorin I.

```
CNNM1 88  PSPTLNSGENGTGDWAPRLV 107
CNNM2 103 LRVYGQNINNETWSRIAFTE 122
CNNM3 64  LRLFGPGFANSSWSWVAPEG 83
CNNM4 76  LRLYGYSLGNISSNLISFTE 95
                .  * : .  .
```

**Supp. Figure 5. N-glycosylation sequon is conserved in CNNMs**

The location of the predicted N-glycosylation residue is indicated. The glycosylated asparagine is highlighted by a box. Amino-acid conservation is indicated below the alignment.

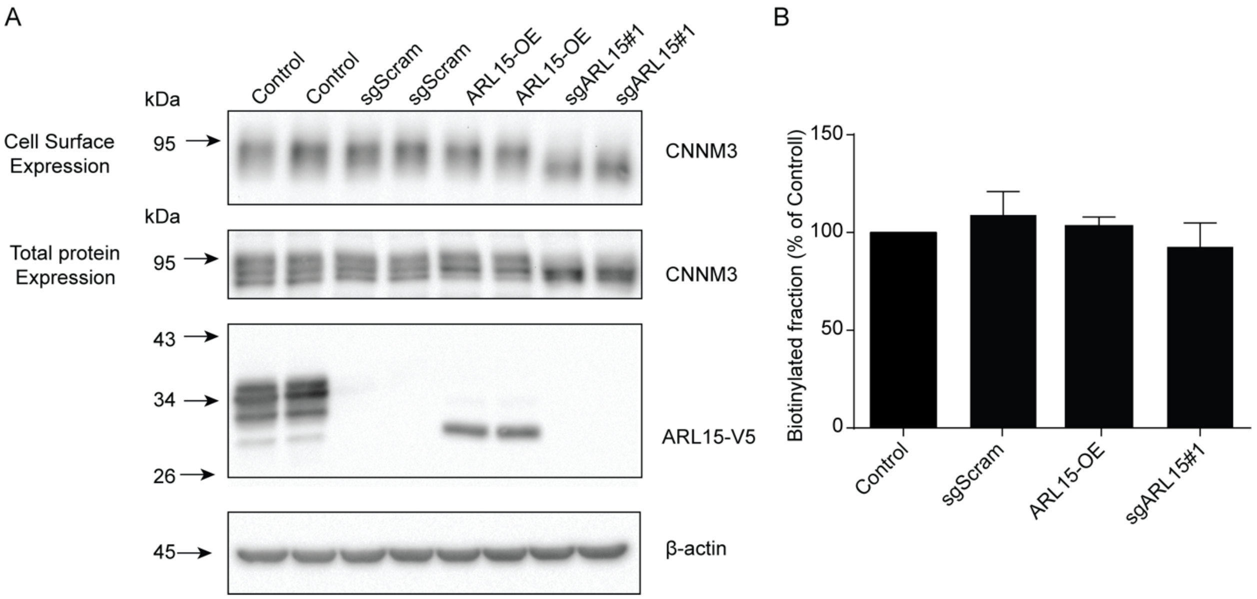


**Supp. Figure 5. ARL15 knockdown does not affect CNNM3 cell surface expression.**

Endogenous cell surface expression of CNNM3 was detected in SK-RC-39 cells by biotinylation. The upper immunoblots show CNNM3 cell membrane expression and the lower blot shows the total CNNM3 expression. (B) The diagram shows the quantification of cell surface CNNM3 expression corrected for total CNNM3 protein expression. Results are the mean ± SEM of 3 independent experiments.

#### Supplementary Tables

#### Supplemental Table 1: Brownian Dynamics data

| No | Size | Repr | ReprE | ClAE | CLAED | ElE | ElDesE | HyDesE | spread | stddev | max |
|---|---|---|---|---|---|---|---|---|---|---|---|
| 1 | 61257 | 141 | -25.858 | -25.609 | 0.743 | -7.640 | 3.766 | -21.984 | 28.666 | 15.109 | 44.108 |
| **2** | **63258** | 277 | **-25.069** | **-26.597** | 1.261 | **-7.960** | **6.939** | **-24.048** | 19.155 | 7.041 | 32.403 |
| 3 | 13185 | 200 | -25.504 | -26.276 | 0.600 | -4.104 | 2.758 | -24.158 | 9.718 | 16.588 | 46.429 |
| 4 | 32649 | 363 | -24.803 | -26.204 | 1.889 | -1.591 | 4.212 | -27.424 | 17.866 | 11.585 | 29.227 |
| 5 | 9200 | 484 | -24.473 | -25.335 | 0.439 | -3.661 | 3.750 | -24.562 | 4.455 | 8.879 | 28.354 |

No: Cluster Number
Size: Number of representative entries for the cluster
Repr: Representative chosen
ReprE: Total interaction energy of the chosen Representative
ClAE: Average total energy of all cluster members weighted with number of representatives.
CLAED: Weighted standard deviation of total energy of cluster entries in the complexes (f55) file
Ele: Electrostatic energy of the representative complex
ElDesE: Electrostatic desolvation energy of the representative complex
HyDesE: Hydrophobic desolvation energy of the representative complex
spread: arithmetic average of rmsd's of each cluster member from the representative, weighted by occupancy
stddev: Stddev of the rmsd's for a given cluster, weighted by occupancy
max: Maximum rmsd within one cluster from the representative
